# Order of arrival and nutrient supply alter outcomes of coinfection with two fungal pathogens

**DOI:** 10.1101/2023.12.20.572661

**Authors:** Elizabeth T. Green, Rita L. Grunberg, Charles E. Mitchell

## Abstract

A pathogen arriving on a host typically encounters a diverse community of microbes that can shape priority effects, other within-host interactions, and infection outcomes. In plants, environmental nutrients can drive tradeoffs between host growth and defense and can mediate interactions between coinfecting pathogens. Nutrients may thus alter the outcome of pathogen priority effects for the host, but this possibility has received little experimental investigation. To disentangle the relationship between nutrient availability and coinfection dynamics, we factorially manipulated nutrient availability and order of arrival of two foliar fungal pathogens (*Rhizoctonia solani* and *Colletotrichum cereale*) on the host grass tall fescue (*Festuca arundinacea*) and tracked disease outcomes. Overall, *C. cereale* infection facilitated infection by *R. solani*, increasing its infection rate regardless of their order of inoculation. Additionally, simultaneous and *C. cereale*-first inoculations decreased plant growth and – in plants that did not receive nutrient addition – increased leaf nitrogen concentrations compared to uninoculated plants. Nutrient addition did not influence infection rates, severity of infection, or plant biomass. These findings highlight the importance of understanding the intricate associations between the order of pathogen arrival, host nutrient availability, and host defense to better predict infection outcomes.

## Introduction

A pathogen arriving on a host typically encounters a diverse community of symbionts and other pathogens that have previously colonized that host [1]. The later-arriving pathogen thus enters a host environment that has been shaped by the earlier-arriving species through competition with other microbes for nutrients and other resources [2] and indirectly priming the host immune system [3]. The effects that an earlier-arriving species has on the success of a later-arriving one are called priority effects [4–6]. They can be either inhibitive [7], when earlier-arriving species make the host uninhabitable for later-arriving species, or facilitative [8], when an earlier-arriving species alters the host environment in a way that makes it easier for later-arriving species to establish [9–11]. The order in which pathogens arrive on a host and the outcomes of those pathogen interactions can shift the infection outcomes resulting in different epidemic trajectories and requiring adaptable interventions [7,11–13].

Pathogens are dependent on their hosts for resources including nutrients. Earlier-arriving species and species arriving on nutrient-rich hosts may not face as much interspecific competitive pressure as pathogens arriving at a host that is already heavily infected or depleted of nutrients [2,14,15]. If the arriving pathogen is a stronger competitor for host resources, they may outcompete the previously arriving species and successfully infect the host. However, if they are not as strong a competitor, they may experience decreased infection success and severity. Once a pathogen has successfully infected the host, the severity of that infection will be dependent on competition pressure from other pathogens and the resource concentration in the host tissue. While attention has focused on priority effects based on competition, priority effects can also be facilitative, particularly when within-host species interactions are mediated by direct or indirect suppression of host defenses.

Environmental nutrients can influence host-pathogen epidemic dynamics by changing host growth and defense strategies [13,16,17]. The nutrients available to the host can affect the host tissue nutrient concentrations that pathogens are feeding on and host allocation to defense or growth [18]. Both growth and defense require resources, so host allocation to defense often comes at the expense of growth [19–21]. Hosts spending resources on defense against early-arriving species may be left vulnerable to attack by later-arriving species [22]. The later-arriving species can exploit this vulnerability and cause more severe disease [10]. The strategy a host employs is dependent on the resources available to it [17,23]. While growth-defense tradeoffs have been well studied in many herbivores and some pathogen systems [3,24] the implications for pathogen coinfection are largely unknown.

Despite the overlapping effects of nutrients on priority effects of pathogens within hosts, and on the growth and defense of the hosts, there has been little experimental work to investigate how infection outcomes are shaped by interactions between order of arrival and nutrient availability. We conducted an experiment that factorially manipulated the nutrient availability to a host grass, tall fescue (*Festuca arundinacea)*, and the order of arrival of two commonly coinfecting fungal pathogens (*Rhizoctonia solani* and *Colletotrichum cereale*) to test three predictions about infection rates and outcomes. First, we predicted that the order of pathogen arrival and nutrient availability will affect infection rates by modulating host susceptibility. Second, we predicted that nutrient availability and pathogen order of arrival will influence the severity of infections by mediating interactions between coinfecting pathogens. Finally, we predicted that coinfection will reduce host growth more under low-nutrient conditions, due in part to resources spent on plant defenses. Understanding the outcomes of our three predictions will help us to understand the dynamics of coinfection under differing environmental nutrients scenarios as humans increasingly alter environmental nutrient supplies [13].

## Methods

### Study System

Tall fescue (*Festuca arundinacea*) is a common grass species that is widely utilized in agriculture and recreation and is also a dominant species in non-cultivated systems across the southern and eastern United States [25]. Tall fescue is host to numerous fungal pathogens, including those infecting both the roots and the leaves. In this study, we focus on two foliar fungal pathogens that frequently coinfect their host. *Rhizoctonia solani* is a necrotrophic parasite that feeds on dead plant tissue. Upon infection, it kills leaf tissue causing necrotic lesions and can ultimately kill the whole leaf. It is soil and water borne and causes the disease brown patch, locally killing clusters of plants [26]. *Colletotrichum cereale* is one of several species responsible for anthracnose disease in tall fescue [27]. It is a hemibiotrophic fungus, initially infecting and extracting nutrients from living cells, then changing its feeding strategy to kill leaf tissue causing necrotic lesions. *C. cereale* has been found to facilitate growth of *R. solani* when arriving first in tall fescue, leading to an increased disease burden [11].

### Experimental Overview

We ran the experiment twice in 2022. For the first temporal replicate, we germinated plants on February 28 and harvested them on May 9 and for the second temporal replicate, we germinated plants on April 11 and harvested them on June 15. We applied two treatments to each plant: a low (50ppm) or high (150ppm) fertilizer treatment (20-20-20 Peter’s Original Water Soluble Fertilizer) and a fungal inoculation (Table 1). In each replicate, the plants were randomly assigned a treatment group and the treatments were factorially applied. We grew the plants in a greenhouse until they were inoculated and then moved them to either growth chambers (replicate 1) or a grow room (replicate 2). The spatial (two growth chambers) and temporal replicates were represented as three blocks in all further analyses. Both locations were kept at 29°C with constant light. While we initially assigned 44 plants to each inoculation group (30 plants in the mock inoculation group), *C. cereale* spore production is highly variable, and we did not have enough inoculum to complete the original plan in the first replicate. We inoculated some of the plants that were originally supposed to receive the *C. cereale* inoculation with *R. solani* only or *R. solani* first treatments after running out of *C. cereale* spores and reduced the number of plants in those groups in the second replicate. In total, there were 244 plants, 6 of which tested negative for the vertically transmitted endophyte *Epichloë coenophiala* and were removed from the data analysis (Table 2).

**Table 1.** Inoculation treatments.

| Inoculation | Day 1 | Day 2 |
| --- | --- | --- |
| Mock | Mock *C. cereale* and mock *R. solani* | NA |
| *R. solani* only | *R. solani* | Mock *C. cereale* |
| *C. cereale* only | *C. cereale* | Mock *R. solani* |
| *R. solani* first | *R. solani* | *C. cereale* |
| *C. cereale* first | *C. cereale* | *R. solani* |
| Simultaneous | *C. cereale* and *R. solani* | NA |

**Table 2.** Number of plants per inoculation group (n = 244 (238 E. *coenophiala* positive and 6 E. *coenophiala* negative)).

| Inoculation | Planned Rep 1 | Rep 1 | Planned Rep 2 | Rep 2 | Total *E. coenophiala* Positive Plants |
| --- | --- | --- | --- | --- | --- |
| Mock | 20 | 20 | 10 | 10 | 30 |
| *R. solani* only | 24 | 31 (1 *E. coenophiala* negative) | 20 | 15 | 45 |
| *C. cereale* only | 24 | 18 (1 *E. coenophiala* negative) | 20 | 20 | 37 |
| *R. solani* first | 24 | 29 (1 *E. coenophiala* negative) | 20 | 15 (1 *E. coenophiala* negative) | 42 |
| *C. cereale* first | 24 | 22 | 20 | 20 (1 *E. coenophiala* negative) | 41 |
| Simultaneous | 24 | 24 | 20 | 20 (1 *E. coenophiala* negative) | 43 |

### Experimental Set-up

Seeds collected from Widener Farm (Duke Forest) in 2018 were primed by soaking in water for 6 hours and allowed to dry overnight. The seeds were sprinkled over moist vermiculate and left in the greenhouse with a 12h light/dark cycle to germinate for 14 days. We haphazardly selected seedlings to transplant one seedling into each pot packed with MetroMix 360 and left them to grow for another two weeks in the greenhouse. We then started applying 10mL of the assigned fertilizer concentration (low or high) twice per week.

The original fungal cultures for both species were collected from Duke Forest in 2015 by F. Halliday and K. O’Keeffe. *R. solani* was grown on potato dextrose agar (PDA) and frozen in potato dextrose broth at −80°C. *C. cereale* was grown on PDA slants and stored at 4°C. Three weeks before inoculation, five plugs of *R. solani* hyphae were pulled from the freezer and plated on plates of PDA. Cultures were left to grow at 25°C with constant light for seven days and then replated to multiply the number of active cultures.

At the same time, five *C. cereale* plugs were pulled from PDA slants in storage and plated on fresh PDA plates. Plates were left to grow for two weeks at 25°C under constant light. We then flooded the *C. cereale* plates with 8mL of water and scraped them with a cell scraper to loosen the spores. The spore solution was collected and 0.5mL of solution was spread on each new plate. The plates were left unparafilmed to stress the cultures and encourage sporulation. After five days each *C. cereale* plate was flooded again with 10mL of sterile water and scraped to release spores. The spore slurry was collected and the spore density from each plate was estimated by looking at the slurry on a hemocytometer under a microscope. The combined spore concentrations were 690,000 condiospores/mL in the first repetition and 568,200 condiospores/mL in the second.

### Inoculations

We randomly assigned plants to one of the six experimental treatments with an equal number between fertilizer treatments (mock *C. cereale* and mock *R. solani,* only *C. cereale*, only *R. solani*, *C. cereale* followed by *R. solani, R. solani* followed by *C. cereale*, and both *R. solani* and *C. cereale* simultaneously; Table 1). The plants that only received one inoculation were treated with a mock version of the other inoculation (potato dextrose broth or a plug of PDA) on the second inoculation day. We haphazardly selected a tiller, shoot growing from the base of the plant, from the one to five tillers of each plant and inoculated the second youngest leaf of that tiller. The tiller was stabilized with a wooden stake and marked by wrapping a single plastic comb binding around the tiller. We first painted the leaf with 0.5mL of the *C. cereale* spore solution or mock potato dextrose broth solution, if receiving that treatment. We then gently wrapped the leaf in tinfoil at the base of the leaf, leaving a gap between the tiller and leaf. With a cork borer sterilized with ethanol and run through flame, we pulled a plug from the outer edge of either active *R. solani* culture or sterile PDA plate and nestled the plug between the leaf and tiller. A piece of cotton was soaked in sterile water and placed on top of the plug before closing the tinfoil wrap around both the plug and cotton. We finished each plant by wrapping the tinfoil casing with parafilm to keep in the moisture. Finally, plants were placed in dew chambers made of plastic bags spritzed with water and rubber banded at the base of the pot. Plants were moved to growth chambers or a grow room with constant light and temperature of 29°C and placed in tubs of water to bottom water.

After 48 hours, we removed the plastic bag, parafilm, tinfoil, cotton, fungal plug, and stake. Four days after the first round of inoculations, we completed the second round of inoculations for the plants in the co-inoculation treatments or the mock treatment for those receiving a single inoculation.

### Infection Surveys

We surveyed the plants five times, every 2-3 days, in each replicate (Table 3). We examined the inoculated leaf for visible *R. solani* and *C. cereale* lesions and measured the total length of each lesion (Figure S1).

**Table 3.** Experimental timeline.

| Rep 1 Days Post Inoculation | Rep 2 Days Post Inoculation | Activity |
| --- | --- | --- |
| -52 | -49 | Germinate seed |
| -38 | -35 | Plant seedlings |
| -24 | -21 | First fertilizer treatment |
| 0 | 0 | First inoculation day |
| 3 | 3 | First survey |
| 5 | 4 | Second inoculation day |
| 7 | 7 | Second survey |
| 10 | 10 | Third survey |
| 14 | 13 | Fourth survey |
| 18 | 16 | Fifth survey and harvest |

At the end of the surveys, we collected the inoculated leaf from every plant and placed them in paper bags for elemental analysis. We also collected a one cm tiller sample to test) for the fungal endophyte, *Epichloë coenophiala* (Agrinostics, Phytoscreen field tiller endophyte detection kit. The prevalence of *E. coenophiala* was 97.6% in all plants, so we removed the endophyte negative plants from the sample. The rest of the plant was collected and dried to measure aboveground biomass. The dried inoculated leaves were ground in a Retsch mixer ball mill. We weighed out between 1.0mg and 3.0mg of ground leaf tissue and packed it into tin capsules. Samples were shipped to the Stable Isotope Analysis Lab at the University of Georgia for elemental analysis. Leaves were analyzed for total carbon (C) and nitrogen (N) content and their corresponding elemental ratio (C:N).

### Data Analysis

All analyses were completed in R (4.2.1) and linear models (lm, stats) were performed with a fixed effect of experimental block and a full factorial of nutrient and inoculation treatment. We ran Tukey HSD as a pairwise comparison between inoculation groups and nutrients for all linear models (emmeans) [28].

We ran all infection analyses for both *R. solani* and *C. cereale.* We analyzed disease progression for the first four surveys after inoculation, because the sequential inoculations only had four surveys after the second inoculation. To assess the rate of infection in each group, we performed a survival analysis of infection incidences across the survey period using a Cox proportional hazards model of infection incidence that evaluates the effects of nutrients, inoculation group, and their interaction (survival) [29]. To analyze infection severity independently of infection success, treatment effects were analyzed separately for each disease, including only plants symptomatic for that disease (*C. cereale* n = 98, *R. solani* n = 107). For both infection types, we calculated the area under the disease progression stairs (AUDPS) of lesion length across the survey days (epifitter) [30]. AUDPS as a metric of disease severity provides an advantage over other metrics like the area under the disease progression curve because if better accounts for the contribution of the first and last survey measurements [31]. We then log transformed the AUDPS values to minimize heteroskedasticity and performed a linear model of the AUDPS value.

To assess plant growth and nitrogen concentration, we removed plants that were not symptomatic for the expected infection by the end of the survey period (n = 229). We then ran three linear models with aboveground biomass, and log transformed C:N ratio [32] and total N (%) as the dependent variable.

## Results

### Rate of Infection

We conducted a survival analysis with a Cox proportional hazard model to assess infection rates between treatment groups. When plants were inoculated with only *R. solani* their infection rate with *R. solani* was 20% lower when compared to the other inoculation groups (p = 0.008, Table S1A, Figure 1A). Generally, *C. cereale* inoculation facilitated infection by *R. solani* regardless of nutrient treatment or order of arrival (ANOVA; inoculation: Chi^2^ _1,146_= 26.19, p < 0.001; nutrients: Chi^2^_3,146_ = 0.74, p = 0.39; inoculation * nutrients: Chi^2^_3,146_ = 1.42, p = 0.70, Table S1B). However, neither *R. solani* inoculation treatment nor nutrients affected the rate of plant infection by *C. cereale* (ANOVA; inoculation: Chi^2^_3,141_ = 0.88, p = 0.83; nutrients: Chi^2^_1,141_ = 1.51, p = 0.22; inoculation * nutrients: Chi^2^_3,146_ = 1.61, p = 0.66, Table S2, Figure 1B).

**Figure 1.**
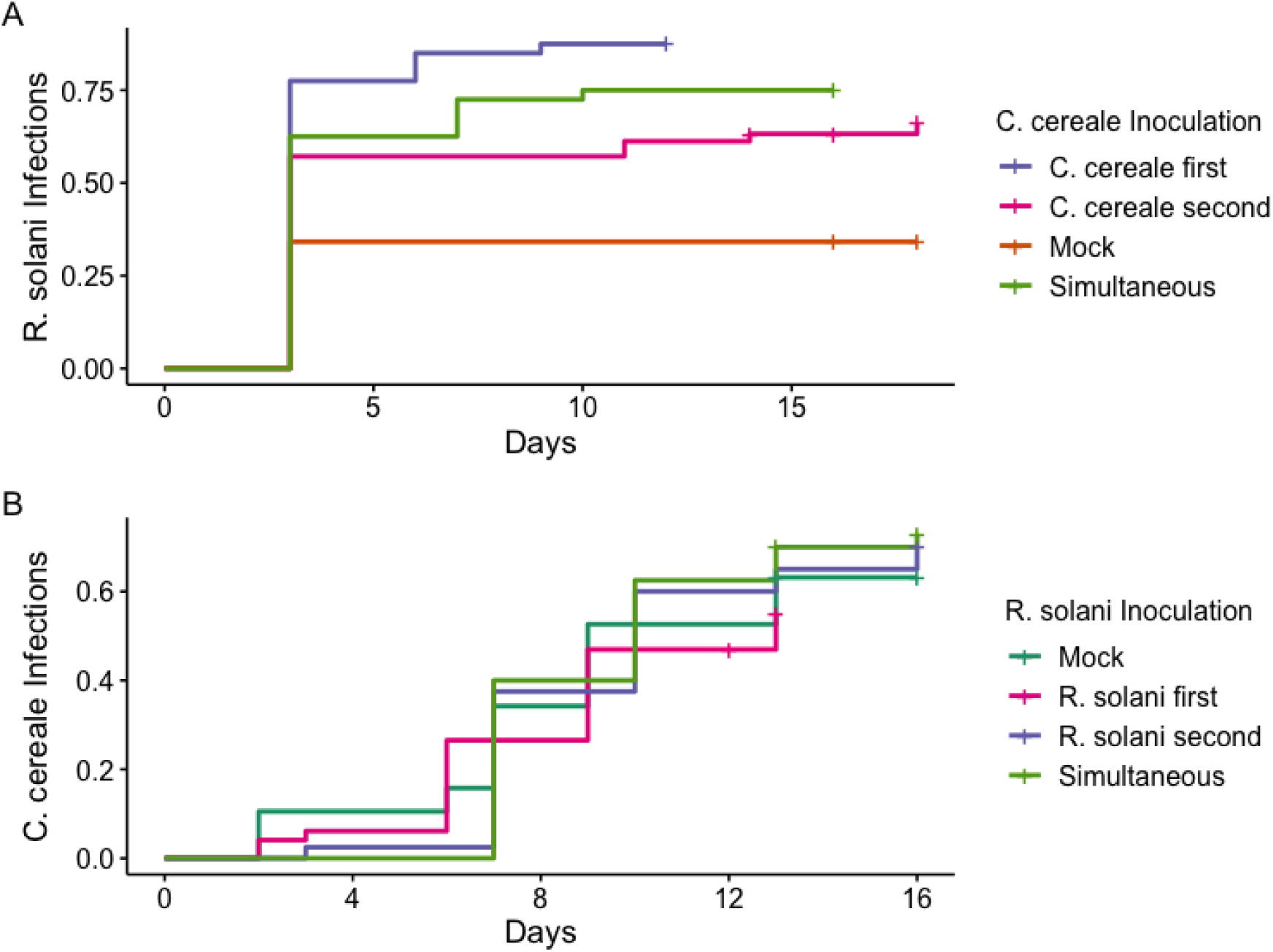
Kaplan-Meier curves of *R. solani* (A) and *C. cereale* (B) infection incidence by nutrient and inoculation treatment. Across nutrient treatments, *R. solani* infection rate when mock inoculated with *C. cereale* (orange) was significantly lower than the other treatment groups.

### Disease Severity

Analyzing the disease progression over time, we found that the severity of *R. solani* lesions in terms of AUDPS were marginally significantly affected by *C. cereale* inoculation (F_3,95_ = 2.52, p = 0.0627, Figure 2A, Table S3).When broken down into pairwise comparisons, plants inoculated with *C. cereale* second experienced 15% more severe *R. solani* lesions than those that were first inoculated with *C. cereale* (Tukey HSD, p = 0.033). We found no effect of fertilization on *R. solani* infection (nutrients: F_1,95_ = 0.01, p = 0.964; inoculation * nutrients: F_3,95_ = 0.88, p = 0.456, Table S3). The severity of *C. cereale* infection in terms of AUDPS was not significantly affected by inoculation group or nutrient treatment (inoculation: F_3,80_ = 1.39, p = 0.251; nutrients: F_1,80_ = 0.24, p = 0.6253; inoculation * nutrients: F_3,80_ = 1.66, p = 0.181, Figure 2B, Table S4).

**Figure 2.**
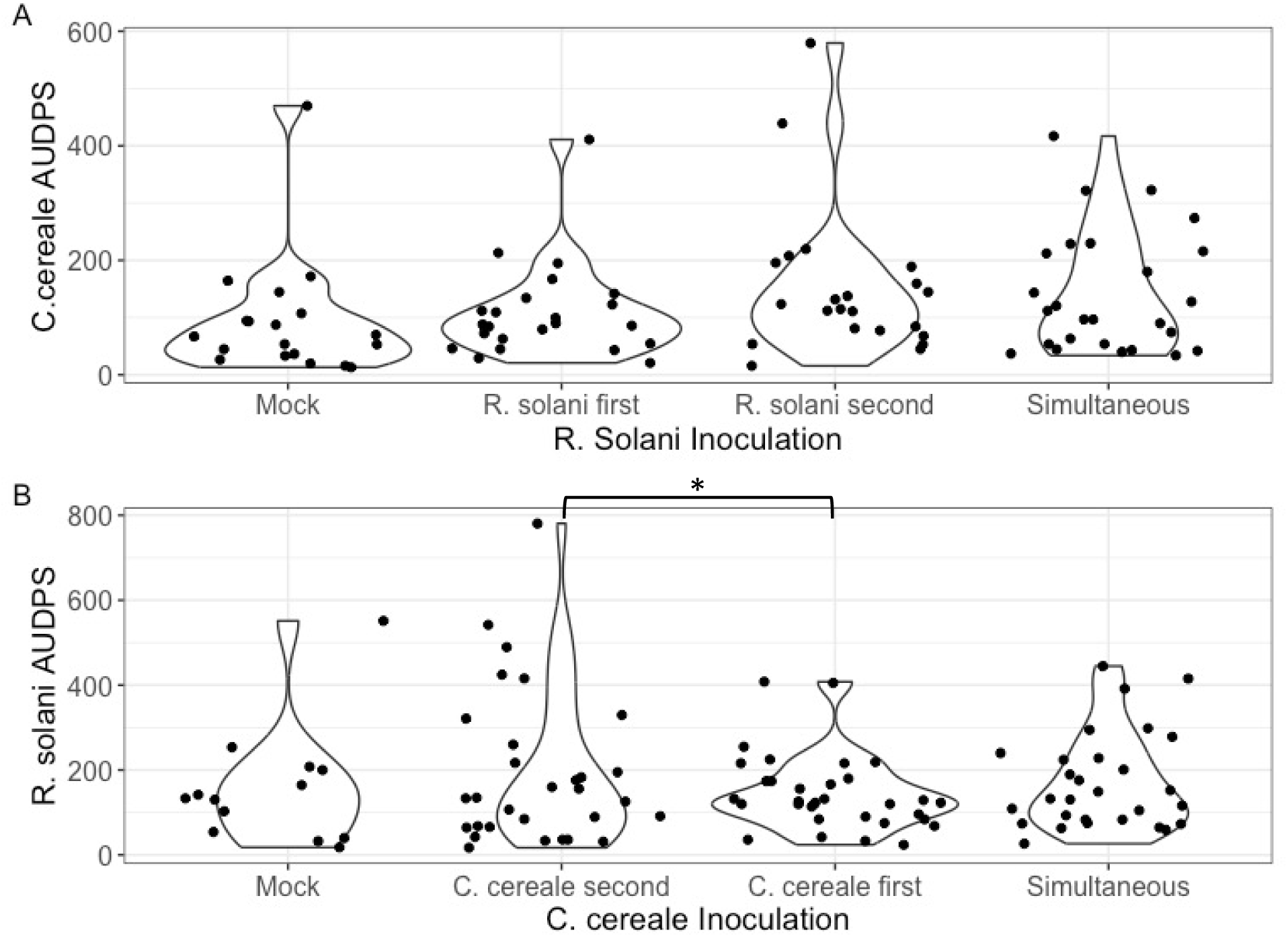
Distribution of area under the disease progress stairs (AUDPS) of *R. solani* (A) and *C. cereale* (B) lesion lengths for plants with symptomatic infection. Significant differences in *R. solani* disease severity between inoculation groups are noted by asterisks.

### Host Growth and Nutrient Allocation

Aboveground biomass differed between inoculation groups (F_5,105_ = 12.15, p < 0.001, Figure 3, Table S5), but not between nutrient groups (nutrients: F_1,103_ = 0.83, p = 0.37; nutrients*inoculation: F_3,103_ = 0.30, p = 0.91, Figure 3, Table S5). Plants inoculated with *C. cereale* first followed by *R. solani* or simultaneously with both pathogens had 16% and 24% reduced biomass respectively compared to the mock inoculated plants (Tukey HSD: *C. cereale* first, p = 0.0019: simultaneous, p < 0.001).

**Figure 3.**
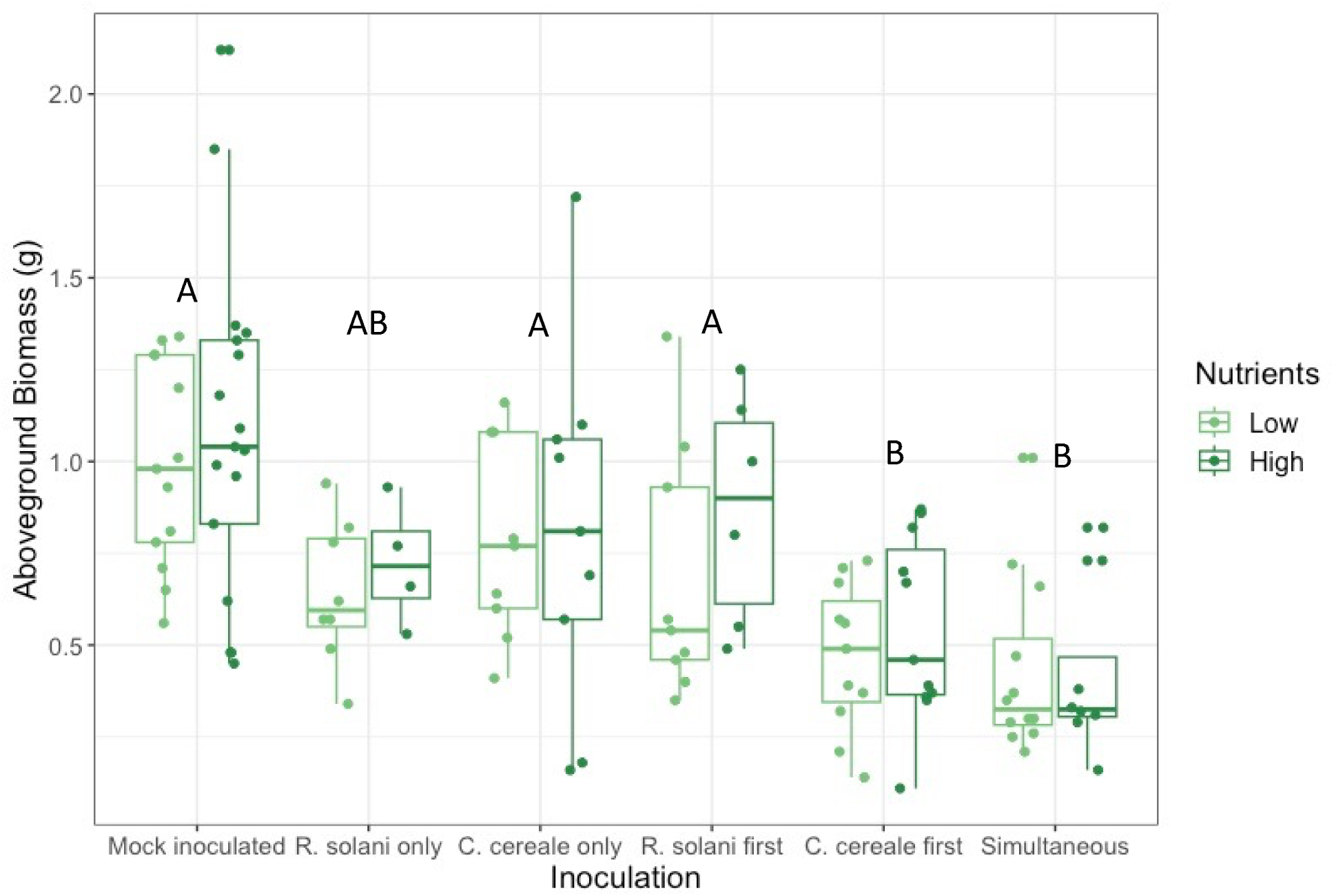
Aboveground biomass of plants by inoculation and nutrient treatment. Box illustrates median and the first (lower boundary) and third (upper boundary) quartile of data with whiskers extending 1.5 times the interquartile range. Significant differences between inoculation groups are noted by letters.

Plants that received high nutrient addition had lower leaf C:N ratios compared to the low nutrient plants regardless of inoculation treatment (nutrients: F_1,86_ = 73.85, p = < 0.001, Figure 4A, Table S6). Of the low nutrient plants, those that were inoculated with only *R. solani* or *C. cereale* had the highest C:N ratio. Mock inoculated plants had 30% lower C:N ratio and plants inoculated with *R. solani* first had 33% lower C:N ratio (Tukey HSD: mock inoculated, p = 0.0306; *R. solani* first, p = 0.0044). Finally, low nutrient plants that were inoculated first with *C. cereale* or with both pathogens simultaneously had at least a 49% reduction in leaf C:N ratios (Tukey HSD: *C. cereale* first, p < 0.001; simultaneous, p < 0.001) and their C:N ratios were similar to all plants in the high nutrient group (inoculation: F_5,86_ = 11.16, p = < 0.001; inoculation * nutrients: F_5,86_ = 5.47, p = < 0.001; Figure 4A, Table S6). Total leaf nitrogen followed similar trends, with the high fertilizer plants having increased leaf nitrogen (nutrients: F_1,83_ = 71.26, p < 0.001, Figure 4B, Table S7) and the leaf nitrogen in low fertilizer plants increasing with simultaneous inoculation or *C. cereale* first inoculation (inoculation: F_5,83_ = 10.55, p < 0.001; inoculation*nutrients: F_5,83_ = 5.05, p < 0.001, Figure 4B, Table S7).

**Figure 4.**
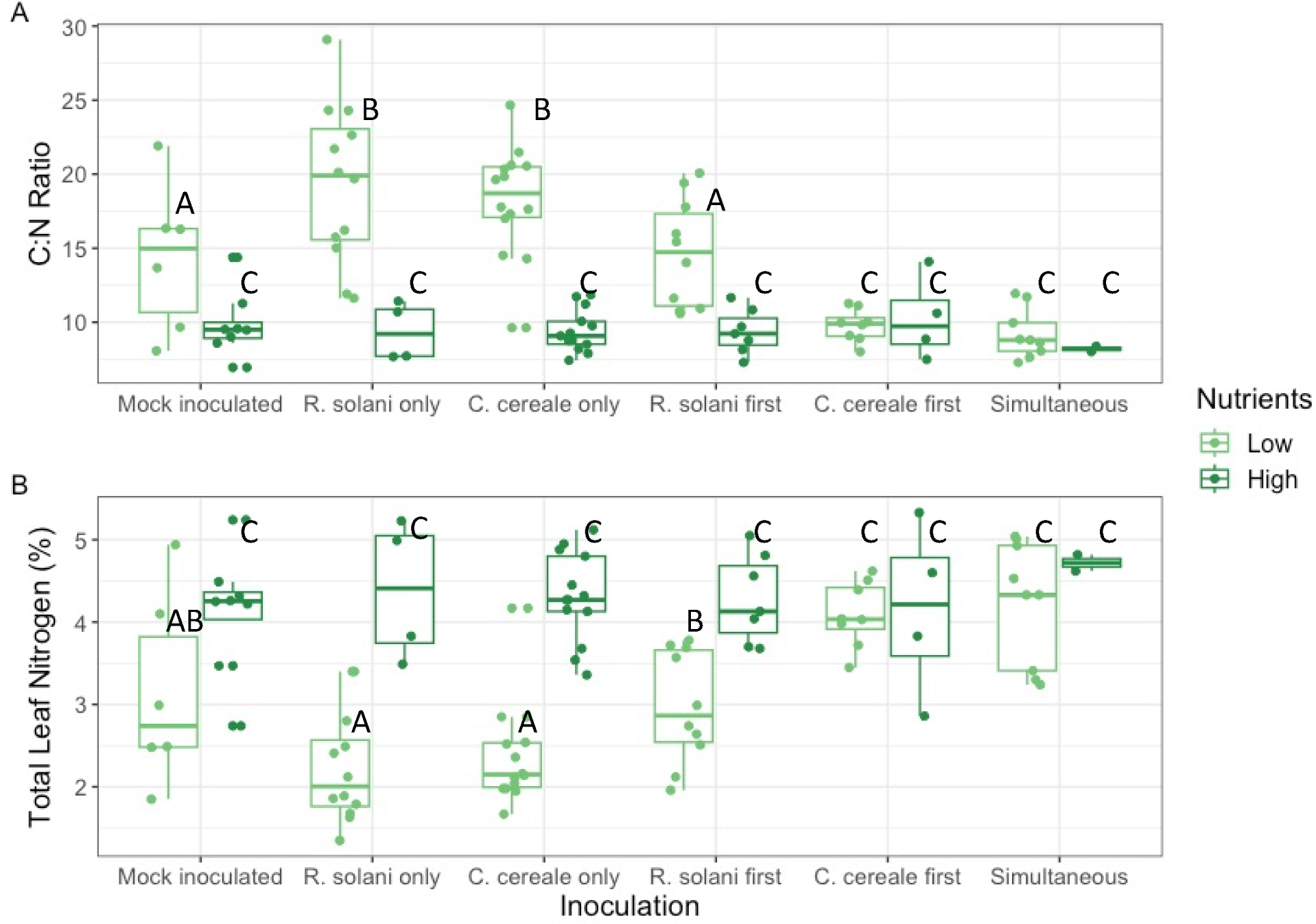
C:N (carbon-to-nitrogen) ratio (A) and total leaf nitrogen (%) (B) of the inoculated leaf by infection type and nutrient treatment. Box illustrates median and the first (lower boundary) and third (upper boundary) quartile of data with whiskers extending 1.5 times the interquartile range. Significant differences between nutrient and inoculation group combinations are noted by letters.

## Discussion

The order of pathogen arrival was the primary force driving infection rate and severity. *C. cereale* facilitated infection rates by *R. solani*. *R. solani* had the lowest infection rate when infecting alone. *C. cereale* also facilitated lesion severity of *R. solani* when arriving second. This facilitation supports our first prediction that the order of arrival would affect infection success by way of the plant defense mechanisms, thereby modulating host susceptibility, and may be explained by interactions between the induced plant defense hormones [4,12,15]. As a hemibiotroph, the primary defense hormone against *C. cereale* is salicylic acid (SA) [22,33]. However, the primary defense hormone against necrotrophs like *R. solani* is jasmonic acid (JA) [22]. When SA is induced, it can inhibit the JA pathway. This interaction has been shown experimentally in many systems [34,35]. By ramping up SA production in response to the arrival of *C. cereale*, the leaf may have been left vulnerable to infection by the necrotophic *R. solani*. Further experiments into the changes in the host caused by *C. cereale* infection are necessary to fully test this mechanism.

Neither pathogen was affected by nutrient availability. Pathogen infection rates and severity were the same between high and low nutrient plants. The lack of interactive effects of nutrient and order of arrival on infection rate or severity suggest that effects of order of arrival on infection rate and severity were not mediated by pathogen competition for nutrients within the leaves. We thus do not have support for our second prediction, that host nutrient availability will influence the severity of infection by mediating competition between coinfecting pathogens. The two focal pathogens occupy different trophic niches within the leaf environment as a necrotroph and a hemibiotroph, so this result indirectly lends support to the niche-based models of historical contingency [5,36].

Aboveground biomass mass not sensitive to nutrient treatment. However, plants receiving the high nutrient treatment had reduced leaf C:N ratios and increased leaf nitrogen concentrations relative to those that received the low nutrient treatment. Infection by both *C. cereale* and *R. solani* simultaneously or with *C. cereale* first reduced plant growth and increased leaf nitrogen in the low fertilizer plants resulting in a decreased C:N ratio similar to that of the high nutrient plants. These results partially support our final prediction. Nutrient availability did not affect plant growth, but it did influence leaf nitrogen concentration and coinfection independently decreased plant growth.

In response to pathogen infection, plants may either divert nitrogen to the damaged leaves to fuel their defense or reduce leaf growth of the damaged area and send carbon to the healthy portion of the plant [13,37]. Total leaf nitrogen (%) of plants in the low fertilizer group increased when plants received two inoculations in any order, supporting that plants were increasing leaf nitrogen to fuel defense when experiencing pathogen coinfections. When the low fertilizer plants received only one inoculation, they had increased leaf C:N ratios and trended towards lower leaf nitrogen, supporting plants pursuing growth when experiencing a single pathogen infection. There is also documented crosstalk between induced defense hormones and growth hormones, suggesting that JA and SA production is at the expense of growth [38,39]. These two mechanisms, increasing leaf nitrogen and decreasing growth under coinfection relative to single infection, support a growth-defense tradeoff. A host’s allocation to growth vs. defense may be critically determined by whether it experiences a single pathogen infection or a coinfection [19–21,40].

Further study of the JA and SA immune pathways is necessary to continue to tease out these priority effects and the nutrient availability necessary for successful defense. This study supports that the interactive effects of nutrients and priority effects are acting on both pathogen infection success and infection severity, but the documented trends do not support competition for nutrients as a mechanism for pathogen priority effects. A better understanding of the mechanisms acting on both infection and severity is important for agricultural and disease management and encourages a closer look at the plant and leaf defense and infection pathways.

## Acknowledgements

This work was supported by the NSF-USDA joint program in Ecology and Evolution of Infectious Diseases (USDA-NIFA AFRI grant no. 2016-67013-25762 and NSF grant DEB-2308472). We would like to thank M. Stroud and N. Harper for their help with inoculations, disease surveys, and sample processing.

