## Supplemental Materials for "Order of arrival and nutrient supply alter outcomes of coinfection with two fungal pathogens"

**Model Outputs**

**Table S1.** Cox proportional hazard model

A. = *R. solani* infection events ~ Block + Inoculation * Nutrient

**df** = 146

| **Parameter** | **Coefficient** | **Hazards Ratio** | **95% Confidence Interval** | **p-value** |
| --- | --- | --- | --- | --- |
| GC2 | 0.274 | 1.315 | 0.844-2.048 | 0.226 |
| GR | 0.188 | 1.207 | 0.744-1.959 | 0.446 |
| H | -0.214 | 0.807 | 0.593-1.585 | 0.569 |
| R | -0.918 | 0.399 | 0.085-0.691 | 0.008 |
| R/C | -0.331 | 0.718 | 0.412-1.812 | 0.699 |
| C/R | 0.261 | 1.299 | 0.776-3.379 | 0.200 |
| H:R | -0.498 | 0.608 | 0.163-2.268 | 0.458 |
| H:R/C | 0.185 | 1.204 | 0.438-3.305 | 0.719 |
| H:C/R | 0.220 | 1.246 | 0.464-3.352 | 0.662 |

B. ANOVA

| **Variable** | **df** | **Chisq** | **p-value** |
| --- | --- | --- | --- |
| Block | 2 | 0.83 | 0.66 |
| Inoculation | 3 | 26.19 | < 0.001 |
| Nutrients | 1 | 0.74 | 0.39 |
| Inoculation*Nutrients | 3 | 1.42 | 0.70 |

**Table S2.** Cox proportional hazards model – ANOVA

= *C. cereale* infection events ~ Block + Inoculation * Nutrient

**df** = 141

| **Variable** | **df** | **Chisq** | **p-value** |
| --- | --- | --- | --- |
| Block | 2 | 0.12 | 0.94 |
| Inoculation | 3 | 0.88 | 0.83 |
| Nutrients | 1 | 1.51 | 0.22 |
| Inoculation*Nutrients | 3 | 1.61 | 0.66 |

**Table S3.** Linear model –

= log(*R. solani* AUDPS) ~ Block + Inoculation * Nutrient - ANOVA

| **Variable** | **df** | **Sum Sq** | **Mean Sq** | **F-statistic** | **p-value** |
| --- | --- | --- | --- | --- | --- |
| Block | 2 | 1.8500 | 0.9250 | 9.4126 | 0.0024 |
| Inoculation | 3 | 0.7425 | 0.2475 | 2.5186 | 0.0237 |
| Nutrients | 1 | 0.0002 | 0.0002 | 0.0020 | 0.6017 |
| Inoculation*Nutrients | 3 | 0.2588 | 0.0863 | 0.8778 | 0.4245 |
| Residuals | 95 | 9.2261 | 0.0983 |  |  |

**Table S4.** Linear model –

= log(*C. cereale* AUDPS) ~ Block + Inoculation * Nutrient - ANOVA

| **Variable** | **df** | **Sum Sq** | **Mean Sq** | **F-statistic** | **p-value** |
| --- | --- | --- | --- | --- | --- |
| Block | 2 | 0.4797 | 0.2398 | 2.1928 | 0.1288 |
| Inoculation | 3 | 0.4767 | 0.1588 | 1.3931 | 0.2510 |
| Nutrients | 1 | 0.0274 | 0.0274 | 0.2403 | 0.6253 |
| Inoculation*Nutrients | 3 | 0.5702 | 0.1901 | 1.6664 | 0.1809 |
| Residuals | 80 | 9.1242 | 0.1141 |  |  |

**Table S5.** Linear model –

= Biomass ~ Block + Inoculation * Nutrients – ANOVA

| **Variable** | **df** | **Sum Sq** | **Mean Sq** | **F-statistic** | **p-value** |
| --- | --- | --- | --- | --- | --- |
| Block | 2 | 0.14 | 0.072 | 2.11 | 0.13 |
| Inoculation | 5 | 2.09 | 0.417 | 12.15 | < 0.001 |
| Nutrients | 1 | 0.03 | 0.028 | 0.83 | 0.37 |
| Inoculation*Nutrients | 5 | 0.05 | 0.010 | 0.30 | 0.91 |
| Residuals | 103 | 3.54 | 0.034 |  |  |

**Table S6.** Linear model –

= log(C:N ratio) ~ Block + Inoculation * Nutrients – ANOVA

| **Variable** | **df** | **Sum Sq** | **Mean Sq** | **F-statistic** | **p-value** |
| --- | --- | --- | --- | --- | --- |
| Block | 2 | 0.85 | 0.40 | 8.64 | < 0.001 |
| Inoculation | 5 | 2.74 | 0.55 | 11.16 | < 0.001 |
| Nutrients | 1 | 3.62 | 3.62 | 73.85 | < 0.001 |
| Inoculation*Nutrients | 5 | 1.34 | 0.27 | 5.47 | < 0.001 |
| Residuals | 83 | 4.07 | 0.05 |  |  |

**Table S7.** Linear model –

~ log(N) ~ Block + Inoculation * Nutrients – ANOVA

| **Variable** | **df** | **Sum Sq** | **Mean Sq** | **F-statistic** | **p-value** |
| --- | --- | --- | --- | --- | --- |
| Block | 2 | 0.78 | 0.39 | 8.54 | < 0.001 |
| Inoculation | 5 | 2.42 | 0.48 | 10.55 | < 0.001 |
| Nutrients | 1 | 3.27 | 3.27 | 71.26 | < 0.001 |
| Inoculation*Nutrients | 5 | 1.16 | 0.23 | 5.05 | < 0.001 |
| Residuals | 83 | 3.80 | 0.05 |  |  |

A)
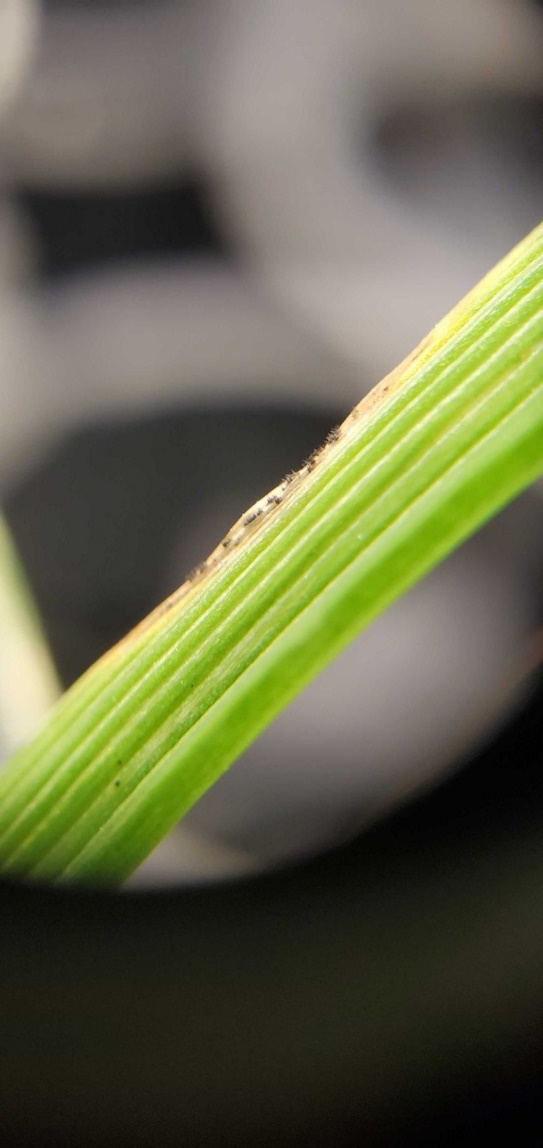
 **B)**
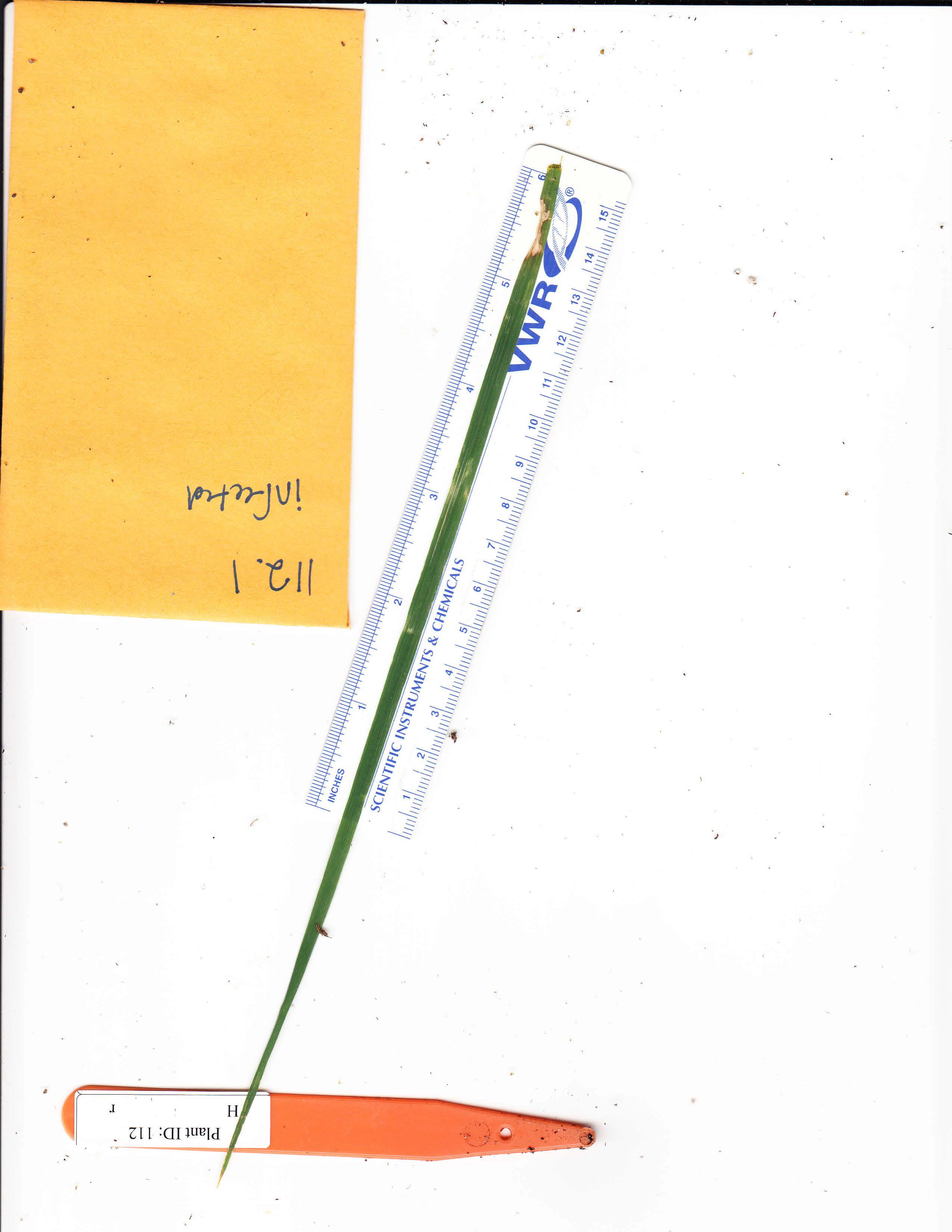

**Figure S1.** Example of A) *C. cereale* and B) *R. solani* lesions.
